# The efficiency for recombineering is dependent on the source of the phage recombinase function unit

**DOI:** 10.1101/745448

**Authors:** Yizhao Chang, Qian Wang, Tianyuan Su, Qingsheng Qi

## Abstract

Phage recombinase function units (PRFUs) such as lambda-Red or Rac RecET have been proven to be powerful genetic tools in the recombineering of *Escherichia coli*. Studies have focused on developing such systems in other bacteria as it is believed that these PRFUs have limited efficiency in distant species. However, how the species evolution distance relates to the efficiency of recombineering remains unclear. Here, we present a thorough study of PRFUs to find features that might be related to the efficiency of PRFUs for recombineering. We first identified 59 unique sets of PRFUs in the genus *Corynebacterium* and classified them based on their sequence as well as secondary structure similarities. Then both PRFUs from this genus and other bacteria were chosen for experiment based on sequential and secondary structure similarity as well as species distance. These PRFUs were compared for their ability in mediating recombineering with oligo or double-stranded DNA substrates in *Corynebacterium glutamicum*. We demonstrate that the source of the PRFU is more critical than species distance for the efficiency of recombineering. Our work will provide new ideas for efficient recombineering using PRFUs.

**Importance:** Recombineering using phage recombinase function units (PRFUs) such as lambda-Red or Rac RecET has gained success in *Escherichia coli*, while efforts applying these systems in other bacteria were limited by the efficiency. It is believed that the species distance may be a major reason for the low efficiency. In this study, however, we showed that it is the source of PRFU rather than the species distance that matters for the recombineering in *Corynebacterium glutamicum*. Besides, we also showed that the lower transformation efficiency in other bacteria compared to that of *E. coli* could be a major reason for the low performance of heterogeneously expressed RecET. These findings will be helpful for the recombineering using PRFUs.

## Introduction

Recombineering using a phage recombinase function unit (PRFU), which usually contains one single-stranded annealing protein (SSAP) and one 5’-3’ exonuclease (EXO), is powerful in certain bacteria (1–5). Expression of the SSAP alone can mediate recombineering with single-stranded DNA (ssDNA) substrates, while expression of the whole system can use double-stranded DNA (dsDNA) as substrates for genome fragment editing (1, 2).

Well-known PRFUs such as lambda-Red or Rac RecET have been used for successful recombineering in *Escherichia coli*. The Red system is phage lambda derived, while the RecET system is from the defective prophage Rac of *E. coli* (1). Lambda-Red contains three adjacent genes *gam, bet*, and *exo*, which encode a host nuclease inhibiting the protein Gam, an SSAP Beta, and an EXO, respectively (6). In *E. coli*, it is possible to mediate recombineering using a dsDNA fragment with a homology arm as short as 35 base pairs through the expression of Red-EXO, Red-Beta, and Red-Gam (7). RecET, in contrast, contains only one SSAP RecT and one exonuclease RecE and has been shown to be highly efficient in linear-linear recombination than lambda-Red (2, 8). Both lambda-Red and Rac RecET function by specifically interacting with their respective partners, except for Red-Gam, which is not necessary for dsDNA-mediated recombineering but can improve efficiency (1, 9).

Previous studies have shown that PRFU functioned well in related species but performed poorly in distant species, presumably because the exonuclease is more species-specific and could not function well heterogeneously (1, 6). To solve this problem, the PRFUs from closely related species were often searched and developed for specific genetic tools (1, 4, 6, 7). Typically, analogues of Beta or RecT were often searched, and EXOs besides these SSAPs were also compared (10, 11). But how the species evolution distance is related to the efficiency of PRFU remains largely unknown, which makes it difficult to mine efficient PRFUs for recombineering.

Here, we present a thorough study of PRFUs in the genomes of the genus *Corynebacterium* to find features that might be related to the efficiency of PRFUs for recombineering. A database to database searching method was first developed to facilitate accurate prediction of novel PRFUs in the genus. Then the identified PRFUs were classified based on both sequential and secondary structure similarity. Typical PRFUs with different sequential and secondary structure similarity as well as host distance to *C. glutamicum* were compared for their ability in mediating recombineering using ssDNA or linear dsDNA substrates. Finally, the recombineering efficiency of the functional PRFUs with their component protein sequence similarity and secondary structure, species relationship, and local genome context were analyzed.

## Results

### Identification of novel PRFUs in the genus *Corynebacterium*

To study the relation of species distance and recombineering efficiency, we first identified the PRFUs in the genus *Corynebacterium*, a taxonomic group with highly diversified species that are of medical, veterinary, or biotechnological relevance (12). Previous studies relating to PRFUs used either reference genomes or nonredundant protein databases for bioinformatics analysis (13–15). As we focused on the relation of species distance and recombineering efficiency, all the bacteria genomes as well as the phage genomes of the genus *Corynebacterium* in the NCBI genome database were used for study.

To facilitate the searching process, a database to database method was first developed (Fig. 1). As PRFU is mainly composed of one SSAP and one EXO, which usually appear next to each other, we studied the existence of both among genomes to improve the prediction accuracy. The prophage origin of candidate PPFUs was also verified by showing their location to the prophage region among genomes (16).

**Fig. 1.**
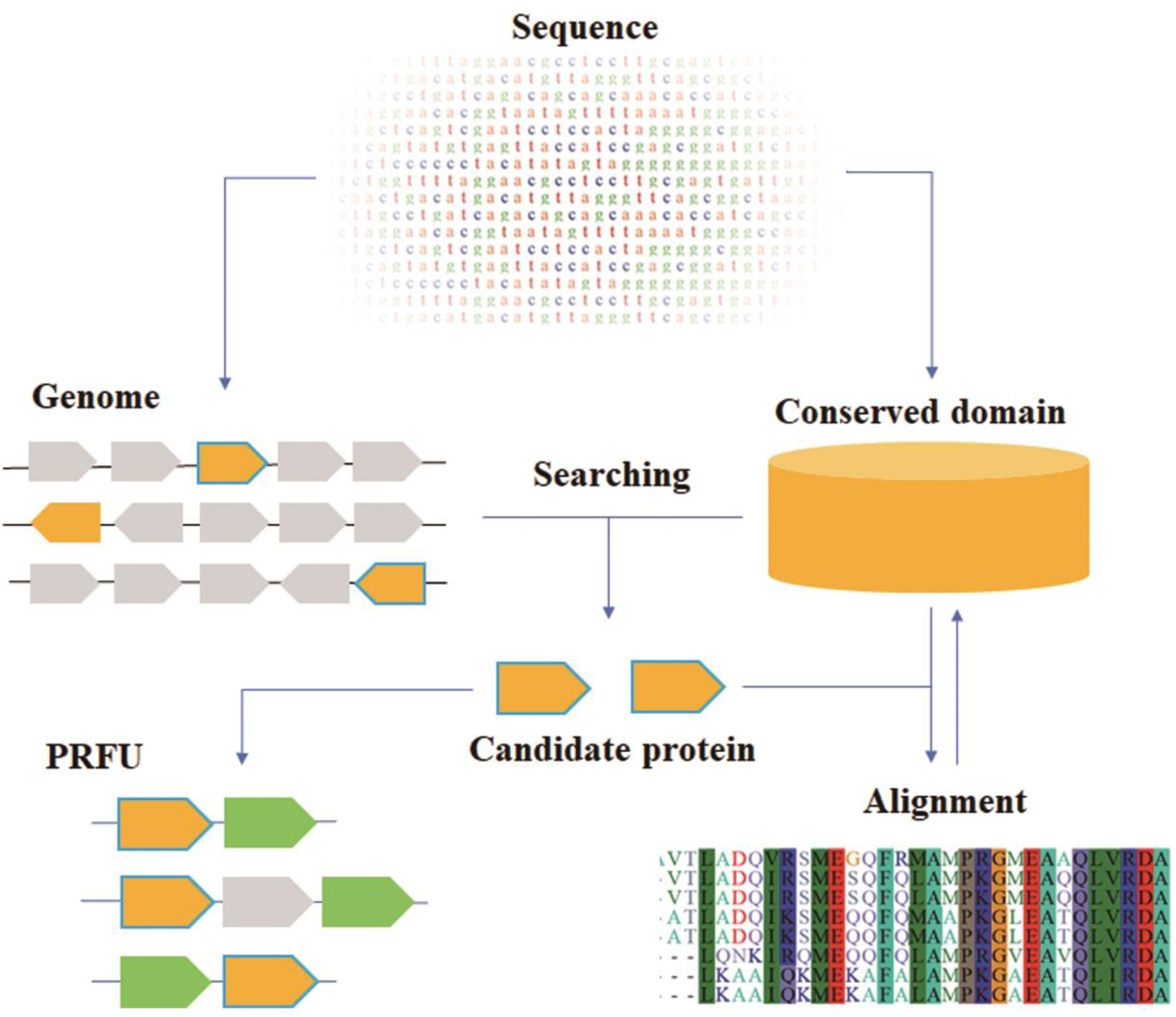
Workflow of finding phage recombinase function units (PRFUs). The genome protein sequence and the conserved domain database sequence were first downloaded from NCBI (https://www.ncbi.nlm.nih.gov/). The genome protein sequence database was then searched in the conserved domain database using the PSI-BLAST method. Candidate single-stranded annealing proteins (SSAPs) or EXOs were then aligned in the conserved domain database to get the consensus parts and form the new conserved domain database. The searching and alignment step were iterated until no new candidates were found. All the candidate SSAPs and EXOs were then collected and paired as PRFUs.

After iterating for three rounds, we identified candidate PRFUs in 84 of the 429 bacteria genomes and none in the phage genomes studied (Fig. S1a and b). We found that among the 84 bacteria genomes containing PRFUs, 52 of them have PRFUs in the prophage region, while 27 of them have similar proteins coded next to PRFUs as those 52 genomes, and five have no obvious features (Fig. S1c). The PRFU component SSAPs were found in 84 genomes, and EXOs were found in 81 genomes. Among them, 54 of the 59 unique SSAPs and 56 of the 57 unique EXOs that were previously annotated as hypothetical proteins were identified.

Analysis showed that these SSAPs and EXOs form 59 sets of unique PRFUs in over 23 species (Table 1). Most of these species have been reported to have no PRFU systems or only have SSAP (13, 17, 18). Different from other species, *C. argentoratense* and *C. glutamicum* both separately have one set of incomplete PRFU with only SSAP but no EXO. Not all the genomes sequenced in these 23 species have PRFUs. This indicates that the PRFU occurrence is genome (strain) related but not species related.

**Table 1.**
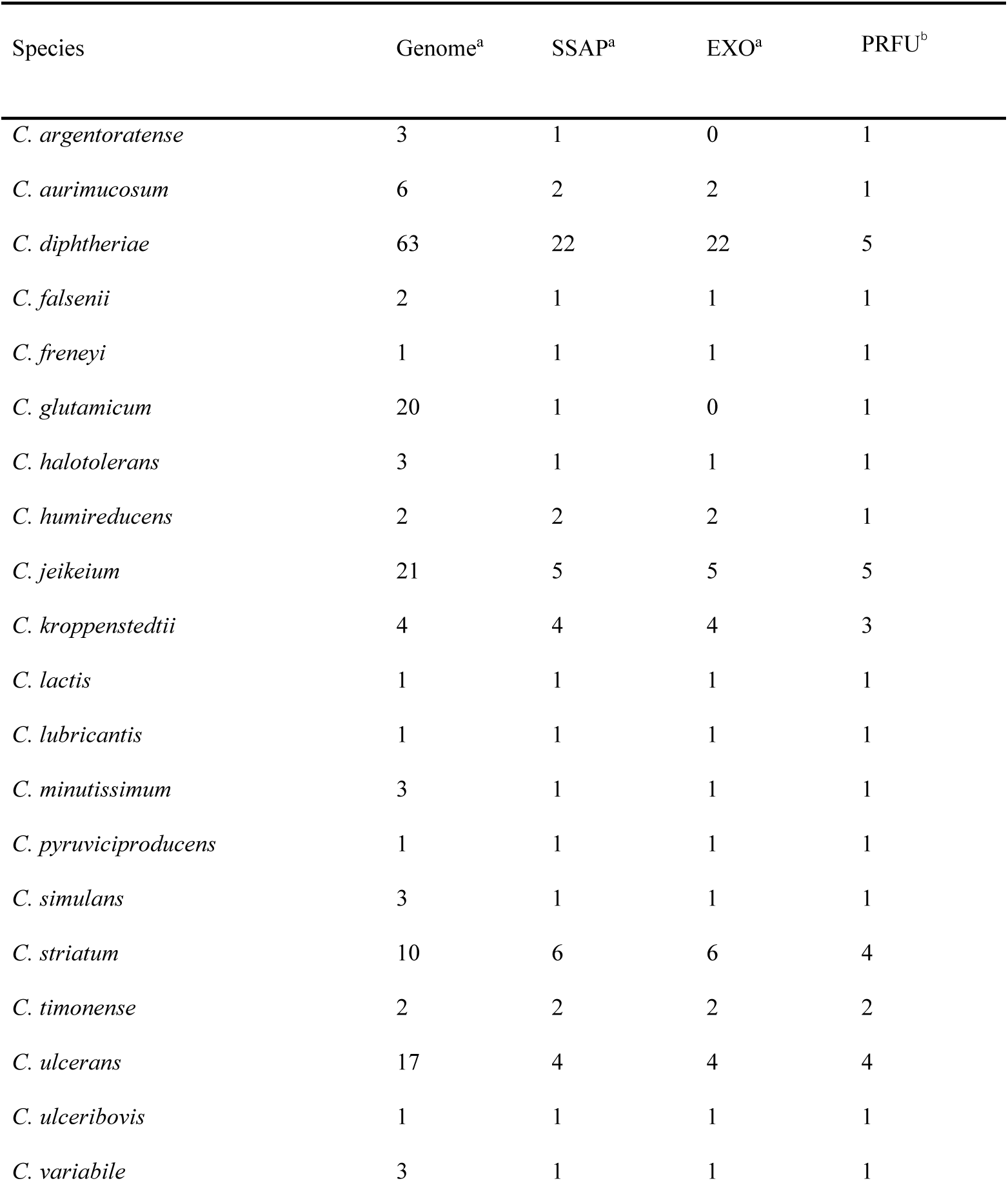
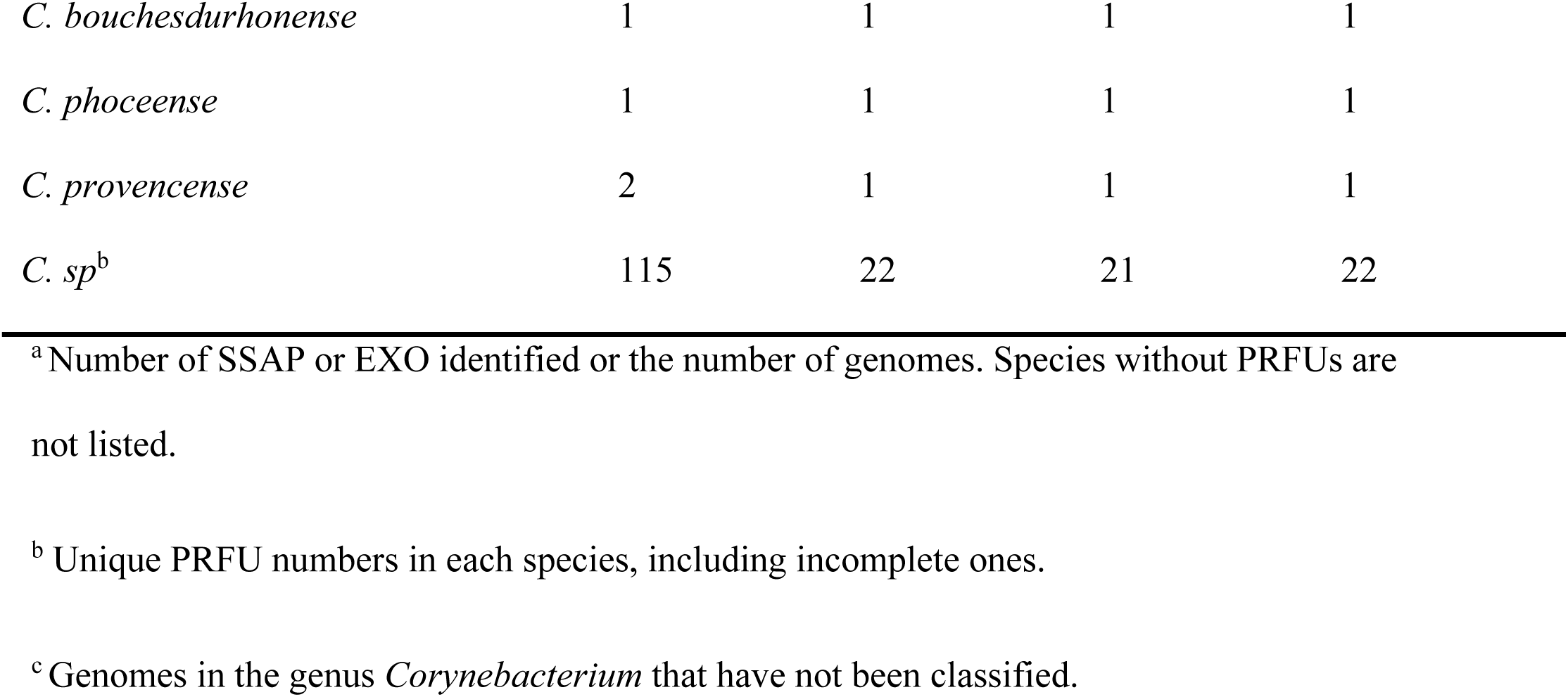
Distribution of PRFUs in the genus *Corynebacterium*.

### Sequence and second structure similarity of the PRFUs and their components

To study the relation of these PRFUs, their component SSAPs and EXOs were classified into five types and three types separately based on both protein sequence similarity and secondary structure similarity (Fig. 2). Each type is conserved in sequence as well as secondary structure and lies within one branch of the phylogenetic trees. Some identified SSAPs or EXOs from other bacteria were also incorporated to construct the tree.

**Fig. 2.**
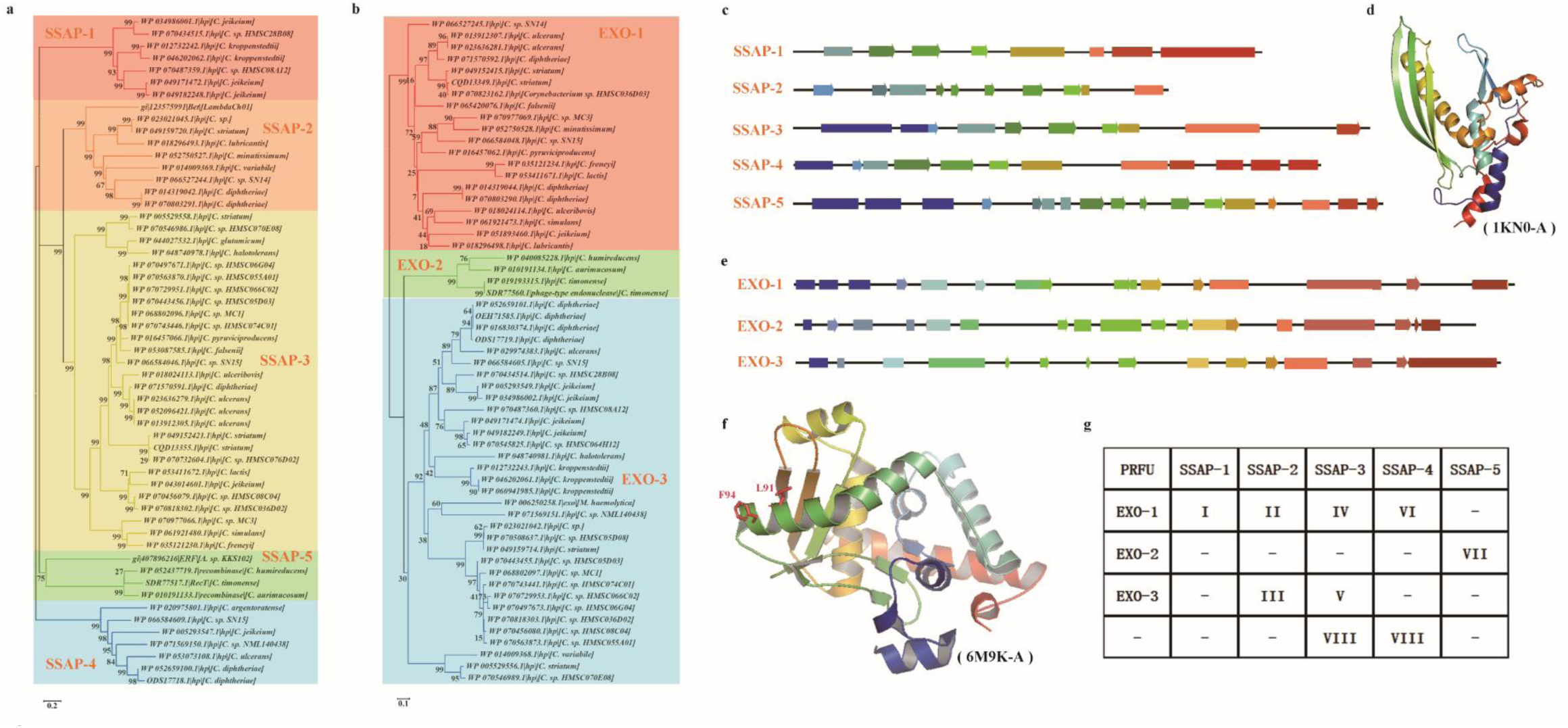
Classification of the PRFUs. **a** Phylogenetic tree and classification of the SSAPs identified. **b** Phylogenetic tree and classification of the EXOs identified. Each type is shown in a different color. The labels are shown as the accession number of the protein, the note for the protein, and the source of the protein. Phylogenetic trees were constructed by MEGA6 using the Neighbor-Joining method. **c** Schematic second structure of each SSAP type. **d** N-terminal structure of human SSAP Rad52. PDB: 1KN0. **e** Schematic second structure of each EXO type. **f** Structure of λ Exo. PDB: 6M9K. **g** Theoretical PRFU type and observed PRFU type. -: not detected.

For SSAPs, the residues in helices are conserved in the same type, while the residues in strands tend to vary from sequence to sequence. Multiple sequence alignment result showed that SSAP-2 shares much similarity to Beta, whereas SSAPs in SSAP-5 were found to have sequence similarity with RecT. A conserved β-β-β-α fold structure was observed in the predicted structure of SSAP-1, SSAP-3, and SSAP-4, while a deformed β-β-β-β-β-α fold structure was observed in SSAP-2 and SSAP-5 (Fig. 2c). The same β-β-β-α fold has been observed in human Rad52, which was found to be composed of highly conserved amino acid residues that are responsible for ssDNA and dsDNA binding (19). The structure of the N-terminus of human Rad52 (Rad52_1-212_) is shown in Fig. 2d as an example. A deformed β-β-α-β-β-β-α fold of Rad52_1-212_ was observed in all SSAP types except SSAP-1. Similar to the Rad52_1-212_, both the C-terminus and the N-terminus of the SSAPs are rich in helices. However, compared to the conserved N-terminus, the residues in the C-terminal region of different SSAP types are irregular. This might be consistent with the fact that the conserved N-terminus of SSAP is responsible for homologous pairing, while the C-terminal interacts with EXO or host factors (20–23).

Unlike SSAPs, the second structure of EXOs varies considerably between types (Fig. 2e). The residues in the N-terminal half of EXOs are conserved across types, while the residues in the C-terminal half vary considerably even in the same type. The residue Leu-91 and residue Phe-94 of λ Exo have been found to be important for forming the λ Exo-Beta complex (23). These residues lie within a helix of the α-α-β-β fold of the λ Exo (Fig. 2f), which was also observed in EXO-3.

These SSAPs and EXOs form eight types of PRFUs in total, which is less than their theoretical combination (Fig. 2g). Typically, SSAP-1 only appears with EXO-1 and forms PRFU-I, while SSAP-5 and EXO-2 only appear with each other and form PRFU-VII, which suggests the conservation of these PRFUs.

### The efficiency of PRFUs for recombineering shows no obvious relation to species distance

Previous studies have indicated that the expression of SSAP alone can promote ssDNA-mediated recombineering, while expression of the full PRFU can facilitate genetic manipulation of the genome with dsDNA (1, 6, 7, 24). To understand the relation of species distance and sequence similarity to recombineering efficiency, PRFUs were chosen based on the species evolutionary relationship as well as the PRFU type. Then both oligo-mediated recombineering and dsDNA-mediated recombineering experiments were conducted in a previously constructed strain C.g-kan (−) (25).

An oligo targeting the defective *kan(-)* was first used for ssDNA-mediated recombineering to recover the function of *kan* (18, 25). As a result, among the eight SSAPs chosen, five of them could promote ssDNA-mediated recombineering in *C. glutamicum* (Table 2, Sap1–Sap8). Then a selection marker interspaced by an 800bp homology arm targeting *recA* was used as the dsDNA substrate to compare the recombineering ability of the full PRFUs whose SSAPs were effective. Though no complete PRFU was found in the *C. glutamicum* R strain, the gene next to its SSAP coding gene was also tested. It was found that complete PRFUs whose SSAP were functional also showed an effect in recombineering, except for the PRFU from *C. variabile*, which was not tested due to the failure in strain culturing (Table 3). The incomplete PRFU from *C. glutamicum* showed no activity in the dsDNA recombineering experiment.

**Table 2.** Recombineering with ssDNA using different SSAPs.

| Strain | Substrate<sup>a</sup> |  |  | Viable cells<br>(x 10<sup>7</sup>)<sup>a</sup> | Transformation<br>efficiency | Recombination<br>efficiency | SSAP type | SSAP source |
| --- | --- | --- | --- | --- | --- | --- | --- | --- |
|  | ssDNA- | ssDNA+ | plasmid (x 10<sup>4</sup>) |  |  |  |  |  |
| Sap0 | 1 ± 1 | 0 | 0.67 ± 0.25 | 41.90 ± 13.11 | 1.36 x 10<sup>-5</sup> | 0 | - | Empty vector |
| Sap1 | 2 ± 3 | 444 ± 40 | 2.13 ± 1.16 | 9.93 ± 3.93 | 2.15 x 10<sup>-4</sup> | 2.08 x 10<sup>-2</sup> | SSAP-5 | *Corynebacterium aurimucosum* DSM 44827 |
| Sap2 | 0 | 61 ± 30 | 3.77 ± 1.93 | 4.03 ± 1.20 | 9.34 x 10<sup>-4</sup> | 1.62 x 10<sup>-3</sup> | SSAP-4 | *Corynebacterium diphtheriae* HC07 |
| Sap3 | 0 | 136 ± 55 | 4.21 ± 0.67 | 16.13 ± 6.38 | 2.61 x 10<sup>-4</sup> | 3.23 x 10<sup>-3</sup> | SSAP-3 | *Corynebacterium glutamicum* R |
| Sap4 | 0 | 14 ± 3 | 2.43 ± 0.41 | 30.63 ± 7.03 | 7.92 x 10<sup>-5</sup> | 5.76 x 10<sup>-4</sup> | SSAP-3 | *Corynebacterium halotolerans* 307 CHAL |
| Sap5 | 0 | 0 | 1.20 ± 0.71 | 18.10 ± 9.92 | 6.65 x 10<sup>-5</sup> | 0 | SSAP-1 | *Corynebacterium kroppenstedtii* DNF00591 |
| Sap6 | 2 ± 2 | 0 | 3.26 ± 1.23 | 41.73 ± 8.27 | 7.82 x 10<sup>-5</sup> | 0 | SSAP-2 | *Corynebacterium lubricantis* DSM 45231 |
| Sap7 | 0 | 0 | 3.26 ± 0.60 | 34.00 ± 9.64 | 9.60 x 10<sup>-5</sup> | 0 | SSAP-2 | *Corynebacterium striatum* 963 CAUR |
| Sap8 | 1 ± 2 | 23 ± 15 | 2.70 ± 0.43 | 16.50 ± 2.83 | 1.64 x 10<sup>-4</sup> | 8.52 x 10<sup>-4</sup> | SSAP-2 | *Corynebacterium variabile* DSM 44702 |
| SmrecT | 1 ± 1 | 20 ± 16 | 1.19 ± 0.30 | 22.07 ± 0.98 | 5.40 x 10<sup>-5</sup> | 1.68 x 10<sup>-3</sup> | SSAP-5 | *Escherichia coli* MG1655 |
| Sap10 | 1 ± 1 | 133 ± 47 | 3.60 ± 0.85 | 0.16 ± 0.04 | 2.30 x 10<sup>-2</sup> | 3.69 x 10<sup>-3</sup> | SSAP-3 | *Corynebacterium sp* HMSC070E08 |
| Sap11 | 0 ± 0 | 5 ± 3 | 8.53 ± 0.92 | 10.67 ± 1.67 | 8.00 x 10<sup>-4</sup> | 5.86 x 10<sup>-5</sup> | SSAP-5 | *Corynebacterium humireducens* DSM 45392 |
| Sap12 | 4 ± 7 | 16 ± 5 | 9.33 ± 1.15 | 10.20 ± 1.40 | 9.15 x 10<sup>-4</sup> | 1.71 x 10<sup>-4</sup> | SSAP-5 | *Corynebacterium timonense* 5401744 |
| Sap13 | - | - | - | - | - | - | SSAP-5 | *Nocardia farcinica* IFM 10152 |
| Sap14 | 0 | 0 | 8.40 ± 0.40 | 9.50 ± 3.25 | 8.84 x 10<sup>-4</sup> | 0 | SSAP-5 | *Proteus mirabilis* HI4320 |
| Sap15 | 0 | 0 | 9.53 ± 0.50 | 9.10 ± 1.74 | 1.05 x 10<sup>-3</sup> | 0 | SSAP-5 | *Aneurinibacillus aneurinilyticus* ATCC 12856 |
| Sap16 | 0 | 0 | 6.67 ± 1.15 | 8.50 ± 1.56 | 7.84 x 10<sup>-4</sup> | 0 | SSAP-5 | *Thermosipho melanesiensis* BI429 |

**Table 3.** Recombineering with dsDNA using different PRFUs.

| Strain | Substrate<sup>a</sup> |  |  | Viable cells<br>(x 10<sup>5</sup>)<sup>a</sup> | Transformation<br>efficiency | Recombination<br>frequency | PRFU<br>type | PRFU source |
| --- | --- | --- | --- | --- | --- | --- | --- | --- |
|  | dsDNA- | dsDNA+ | plasmid (x 10<sup>5</sup>) |  |  |  |  |  |
| ET1 | 0 | 2365 ± 822 | 1.52 ± 0.09 | 3.73 ± 1.44 | 4.08 x 10<sup>-1</sup> | 1.56 x 10<sup>-2</sup> | VII | *Corynebacterium aurimucosum* DSM 44827 |
| ET2 | 0 | 4 ± 2 | 3.11 ± 2.07 | 6.10 ± 1.87 | 5.1 x 10<sup>-1</sup> | 1.29 x 10<sup>-5</sup> | VI | *Corynebacterium diphtheriae* HC07 |
| ET3 | 0 | 0 | 1.53 ± 0.50 | 8.5 ± 1.97 | 1.8 x 10<sup>-1</sup> | 0 | VIII | *Corynebacterium glutamicum* R |
| ET4 | 0 | 85 ± 73 | 0.66 ± 0.23 | 11.43 ± 8.44 | 5.73 x 10<sup>-1</sup> | 1.3 x 10<sup>-3</sup> | IV | *Corynebacterium halotolerans* 307 CHAL |
| ET8 | - | - | - | - | - | - | III | *Corynebacterium variabile* DSM 44702 |
| ET9 | 1 | 2456 ± 391 | 1.02 ± 0.17 | 6.03 ± 1.42 | 1.69 x 10<sup>-1</sup> | 2.41 x 10<sup>-2</sup> | - | *Escherichia coli* MG1655 |

<sup>a</sup> Cell numbers are shown as the average ± S.E.M. Values indicated are the average of three or more experiments. Transformation efficiency was calculated as the number of cells transformed with plasmid of the viable cells. Recombination efficiency was calculated as the recombinants of the successfully transformed cells (cells transformed with plasmid). ET9 expressing RecET was used as the positive control. dsDNA-: no dsDNA control. dsDNA+: addition of 1 µg dsDNA substrate. plasmid: addition of 500 ng plasmid substrate. -: not available.

*C. halotolerans, C. aurimucosum, C. diphtheriae*, and *C. striarum* all have similar distance to *C. glutamicum*, but PRFUs from these species were either functional or nonfunctional, which is also true regarding the PRFUs from *C. variabile* and *C. lubricantis* (Fig. S2, Tables 2 and 3). The efficacy of the functional PRFUs varied regarding their host distance to *C. glutamicum*. Both RecET from the distant *E. coli* and PRFUs from the closely related C. *aurimucosum* showed relative high efficiency in promoting recombineering in *C. glutamicum*, while PRFUs from the closely related *C. halotolerans* and *C. diphtheriae* were more than two order of magnitude less efficient.

### Local genome context is related to the function of PRFUs

It has been proven that the products of genes besides the genes coding SSAP or EXO, such as Red-Gam, may also affect the recombineering efficiency (1, 4, 7). These PRFUs have been shown to be prophage derived, while the evolutionary relation of prophages might be different from their hosts (26). These inspired us to further investigate the relationship of the local genome context and the recombineering ability of PRFUs.

To compare the local genome context of multi-genomes, we considered the five proteins coded upstream and downstream of SSAP. A value S_dis_ of two local genome contexts was calculated to evaluate their distance. As a result, it was found that four of the five functional SSAPs were from genomes sharing a similar local genome context (Fig. 3a). Through the analysis of the distance of the local genome context of other species to that of *C. glutamicum*, it was found that the local genome contexts of these functional ones have obviously smaller and wider distance range (S_dis_ from 0 to 35) than others (S_dis_ from 35 to 40, Fig. 3b).

**Fig. 3.**
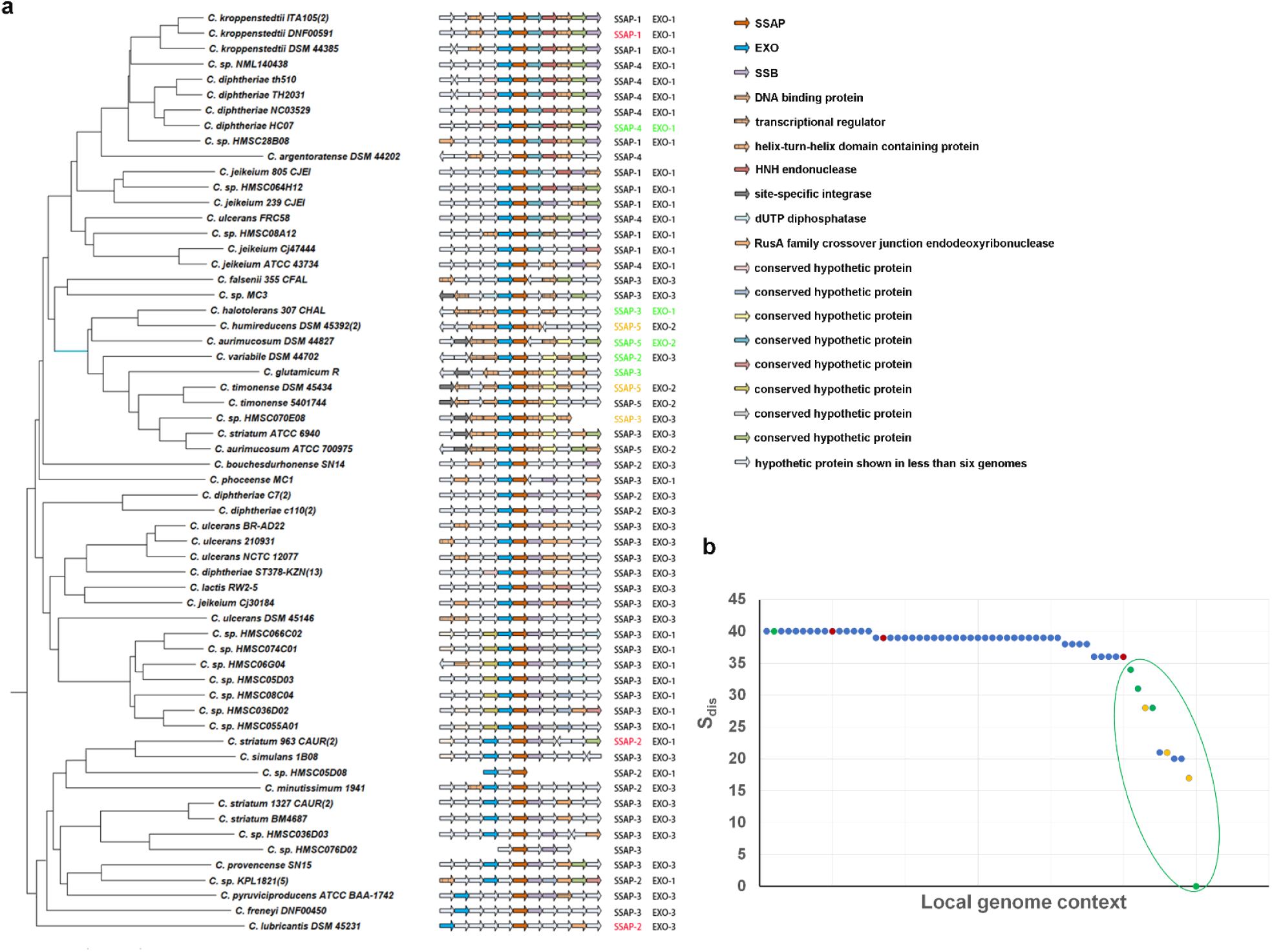
Local genome context of the PRFUs. **a** A phylogenetic tree of local genome contexts of PRFUs. **b** The local genome context distance (S_dis_) of different species with PRFUs compared to that of *C. glutamicum*. PRFUs functioning in the recombineering experiment were colored in green. Those that showed no function were colored in red. Additional verified functional SSAPs were colored in orange.

To test whether the conservation of the local genome context might be related to the recombineering function of PRFUs in *C. glutamicum*, we verified the function of three additional SSAPs whose S_dis_ are smaller than 35 (Table 2, Sap10–Sap12). All the SSAPs in SSAP-5 have S_dis_ less than 35, while RecT and SSAP from *C. aurimucosum* are all highly similar to SSAPs in SSAP-5 and showed high performance in recombineering. Therefore, we also tested four SSAPs from other bacteria who were highly identical to those in SSAP-5 to test whether SSAPs that are highly similar to SSAPs in SSAP-5 may be efficient (Table 2, Sap13–Sap16). As a result, all the selected SSAPs sharing a conserved local genome context were proven to be functional in mediating recombineering, while SSAPs that were highly conserved as SSAP-5 from other bacteria showed no recombineering efficiency. This indicates that the conservation of the local genome contexts might be related to the function of PRFUs.

## Discussion

Although genome sequence analysis has grown rapidly these years, few PRFUs have been identified in the genus *Corynebacterium*. This, together with the phage origin of verified PRFUs, led us to investigate all the genomes sequenced in the genus. As each genome represents a strain record, the occurrence of PRFUs in both reference and non-reference genomes and the irregular distribution pattern of PRFUs among genomes within the same species verifies the idea that the existence of PRFUs is strain related but not species related. When searching for PRFUs among all the plasmids of the genus *Corynebacterium*, only one SSAP coding gene was found on the plasmid of the *C. glutamicum* R strain, which may explain why no PRFU was previously found in *C. glutamicum* when searching its reference genome (13, 18). These indicate that the prediction of PRFUs using reference genomes might lead to errors in conclusion about their existence in the species, while using all the genomes as this study did or using nonredundant protein databases as in other reports may avoid this problem (1, 7, 27).

To compare the ability of PRFUs, we verified several novel identified ones with both ssDNA and linear dsDNA-mediated heterogeneous recombineering in *C. glutamicum*. Some PRFUs tested showed a relatively high ability in conducting recombineering as previously reported tools in other bacteria (1, 6, 7). The recombination efficiency was as high as about 10^−2^, which means there will be one recombinant over one hundred cells successfully transformed. This suggests the great potential of these PRFUs for recombineering in the genus *Corynebacterium*. However, as we have shown in addition to other studies (6, 15), the relative low transformation efficiency could be a major limit of recombineering efficiency in other bacteria other than the PRFUs used.

It has been believed that the efficiency of PRFU for recombineering was related to the species distance, which led to the study of PRFUs in the native species of several bacteria (6, 10, 27). In this study, the host species where most of the tested PRFUs belong to share similar distances to *C. glutamicum*, but the tested SSAPs were either functional or non-functional. RecET from *E. coli* showed relatively high ability in promoting recombineering as those from closely related species of *C. glutamicum* at a recombination efficiency as high as about 10^−2^. The phenomenon that PRFUs from distantly related species performed similarly as those from closely related species (at the same order of magnitude) has also been observed in other reports (1, 7, 18). These suggest an alternative explanation of the relation of the species evolution distance and recombineering efficiency. It was observed in this study that the local genome context conservation in the genus *Corynebacterium* is related to the recombineering function of PRFUs in *C. glutamicum*, but RecET from *E. coli* shows no such conservation. Considering the different performances of PRFUs in this study and previously in *E. coli(1)*, it seems that the efficiency of PRFUs for recombineering is dependent on their source.

In this study, we developed a database to database searching method to facilitate accurate prediction of novel PRFUs in a species that was reported to have no such system. Analysis of the distribution of PRFUs among genomes indicates that their occurrence is related to strains but not species. This is consistent with their features as mobile elements and may provide new insights for the data mining of mobile elements. Through the analysis of the recombineering efficiency of PRFUs with their component protein sequence similarity, second structure, species relationship, as well as local genome context, we demonstrate that the efficiency of PRFUs for recombineering is dependent on the source of the recombinase. This finding will provide new insights for the study of PRFUs for recombineering.

## Methods

### Analysis of PRFU across genomes

Genomes of the genus *Corynebacterium*, including chromosomes, plasmids, and phages were downloaded from NCBI (https://www.ncbi.nlm.nih.gov/) genome database. The conserved domain data of SSAP and EXO from the Conserved Domains Database of NCBI (Accession number: cl04285, cl04500, cl01936, and pfam09588) were also downloaded. Then the following process was iterated.

1. Multiple sequence alignments were done to get the consensus of SSAPs and EXOs and construct database SSAPdb as well as EXOdb.
2. Then genomes of the genus *Corynebacterium* were searched for potential SSAP and EXO in the SSAPdb and EXOdb separately through PSI-BLAST for one iteration.
3. Potential SSAP and EXO were extracted and added to the SSAPdb and EXOdb, respectively.

This process was iterated for three times until no new potential SSAP or EXO was found. Potential SSAPs were chosen from the result of PSI-BLAST using a cut off E value of 10^−6^. Potential EXOs were chosen by setting a cut off E value as 10^−5^.

Prophage was predicted by PHASTER (http://phaster.ca/) (16). The genome location of PRFUs as well as the local genome context were used for the verification of their prophage origin.

### Protein sequence and structure analysis

Multiple protein sequence alignment was performed with MUSCLE (28). The alignment results were then edited manually to get the consensus parts and shown as the default settings in ClustalX (29).

All of the second structures of the alignments were predicated by Jpred4 (http://www.compbio.dundee.ac.uk/jpred4/index.html) (30).

### Alignment of the local genome context

The adjacent five proteins coded upstream and downstream of the SSAP of each genome were extracted to evaluate the local genome context similarity if available. First, the similarity of these proteins was analyzed through PSI-BLAST. Protein pairs with an E value less of than 10^−5^ were grouped together. The groups with proteins of similar annotation or the proteins that were not grouped in the previous step but had similar annotation were merged into one group. The proteins that were not grouped yet were each assigned a group separately.

A similarity matrix of the local genome context was then calculated through the similarity score S_sim_ between each pair of them. The similarity score S_sim_ contained two parts, S_sam_ and S_loc_. S_sam_ was calculated as the sum number of protein pairs from the two genomes that are of same group. S_loc_ was the sum of S_i_, which was calculated in the following way: for each protein coded upstream of SSAP and downstream of SSAP, its position was assigned as “site.” Site i has a range from 1 to 11, corresponding to the position from the farthest upstream position of SSAP to the farthest downstream. If the two proteins of both genomes at the site i belong to the same group, then S_i_ was assigned 1, 2, 3, 4, 5, 6, 5, 4, 3, 2, or 1 for each i from 1 to 11 accordingly. This can be written as

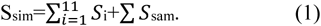

The distance matrix of these genomes was obtained from the similarity matrix, which used a score S_dis_. S_dis_ that was calculated as

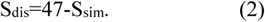

The number 47 was S_sim_ for a pair of completely same local genome context that had an ideal S_dis_ value of 0. As a different site was assigned a different weight, we called these site-weighted scores.

### Phylogenetic analysis

A genome phylogenetic tree was constructed using the 16S rRNA of the representative genomes. A protein phylogenetic tree was constructed using the multiple sequence alignment results of SSAPs and EXOs. For the construction of a local genome context phylogenetic tree, a distance matrix was used as the input. The phylogenetic trees were constructed by MEGA6 using the Neighbor-Joining method with the default settings except for the Interior-branch Test value, which was set as 1000 (31).

### Strain cultivation

The *C. glutamicum* strain C.g-kan (-) was cultured in Luria-Bertani (LB) or brain heart infusion-supplemented (BHIS) medium. For the preparation of the electrocompetent cells, the strains were first cultured in LB medium overnight at 30°C. The cultures were then added to 100 ml BHIS medium at a start optical density of 0.3 and cultured at 30°C until the optical density reached 0.9, with isopropyl β-D-1-thiogalactopyranoside (IPTG) and antibiotics added according to the circumstance. The culture was then chilled on ice for 30 min and centrifuged at 4°C to collect the cells. The sediments were washed four times using 30 ml 10% (v/v) glycerol and resuspended in 1 ml 10% (v/v) glycerol. Then a volume of 100 μl was divided into each tube and stored at −80°C for later experiments. To maintain the plasmids, a final concentration of 100 μg/ml spectinomycin, 15μg/ml kanamycin, or 10 μg/ml chloromycetin was added to the corresponding cultures. For induction purposes, a final concentration of 0.25 mmol/L IPTG was used (25, 32).

### Statistics

The ratio of the number of sequence finished genomes with PRFUs and sequence finished genomes as well as the ratio of the number of sequence unfinished genomes with PRFUs and sequence unfinished genomes were used for a t-test (Table S1).

### Recombineering with PRFUs

The plasmid, ssDNA, and dsDNA substrates were transformed into *C. glutamicum* electrocompetent cells by electroporation. Different substrates were pre-chilled and added to the electrocompetent cells according to the experiment. For recombineering with ssDNA, 2 μg oligo was added to the competent cells. For plasmid and dsDNA, 500 ng or 1 μg substrate was used. The mixture was then used for electroporation at 1.8 kV/mm, 200 Ω, 25 μF. Then 1 ml 46°C BHIS medium was instantly added. The mixture was heat shocked at 46°C for 6 min and then cultured for outgrowth at 30°C, 200 rpm up to 1 h for strain construction purposes or 30°C, 150 rpm for 5 h for the transformation of ssDNA, dsDNA, and control plasmid. Cultures were then plated directly after collecting the cells for most of the time. To count the number of viable cells, 100 μl of 10^−4^ dilution was plated on BHIS with chloromycetin added (15, 18).

Fragments were prepared through overlap extension PCR using TransStart FastPfu Fly DNA Polymerase (TransGen, AP231-03) and collected using a DNA Gel Extraction Kit (TsingKe, GE101-200). Results were verified by agarose gel electrophoresis and Sanger sequencing.

## Acknowledgments

This work was supported by grants from the National Natural Science Foundation of China [31730003, 31670077] and Natural Science Foundation of Shandong Province [ZR2017ZB0210].

We thank Haiying Jin for her help in the recombineering with ssDNA experiment. We also thank Qilong Qin for reviewing the manuscript.

We thank LetPub (www.LetPub.com) for its linguistic assistance during the preparation of this manuscript.

## Author contributions

YZ designed and conducted the experiment, prepared and revised the manuscript. QS designed, supervised the study and revised the manuscript. QW revised the manuscript. TY participated the experiment.

## Supplemental material

**Figure S1. PRFUs in the genus *Corynebacterium*. a** Genomes with or without PRFUs. **b** Genomes in species with or without PRFUs. **c** The phage origin of predicted PRFUs. F: genomes with PRFUs, NF: genomes without PRFUs. CL: PRFUs found in genomes that have been classified into species, UCL: PRFUs found in genomes that have not been classified into any species, F: genomes with PRFUs, NF: genomes without PRFUs. Numbers indicate the genome number.

**Figure S2. Species 16sRNA distance to *C. glutamicum***.

**Table S1. The occurrence of PRFU and genome assemble level.**

**Table S2. Strains and plasmids used in the study.**

